# Fundamental-realized niche contrasts shape multi-scale species coexistence

**DOI:** 10.64898/2026.07.03.736382

**Authors:** Jörn Pagel, Martina Treurnicht, Karen J. Esler, Frank M. Schurr

**Affiliations:** Institute of Landscape and Plant Ecology, University of Hohenheim, Stuttgart, Germany; Centre for Biodiversity and Integrative Taxonomy (KomBioTa), University of Hohenheim & State Museum of Natural History, Stuttgart, Germany; Botanical Society of South Africa, Cape Town, South Africa; Department of Conservation Ecology and Entomology, Stellenbosch University, Stellenbosch, South Africa

## Abstract

Ecological theory states that the geographic ranges and coexistence of species are determined by fundamental and realized niches — the sets of environments where a species’ intrinsic population growth rate is positive in the absence and presence of competitors, respectively. Yet large-scale tests of niche theory have been hampered by the challenge to obtain sufficient data on demography and competition. Here, we quantify fundamental and realized niches by combining data on variation in fundamental demographic rates, community composition and the abiotic environment across the global geographic ranges of 29 shrub species from the South African Fynbos biome (a global biodiversity hotspot). Estimated pairwise competition coefficients and fundamental-realized niche contrasts reveal multi-scale mechanisms of species coexistence. At small scales, species generally exert stronger competition on themselves than on other species. At biogeographical scales, more competitive species have narrower fundamental niches but are not significantly better dispersed, which provides evidence for a generalist-specialist trade-off rather than a competition-colonization trade-off. Under both present and future climates, interspecific competition more strongly limits the realized niches and geographic ranges of generalist species. The large-scale application of niche theory thus identifies key forces shaping biodiversity and indicates that generalist species may be more strongly impacted by climate change than previously thought.

## Introduction

Ecological niches are central for understanding where species can persist and how they interact across environmental gradients. While abiotic environmental conditions determine which plant species can potentially survive, biotic interactions shape coexistence, community composition, and ultimately species’ geographic distributions. Recognizing how these interactions structure plant communities across spatial scales is essential for understanding ecological and evolutionary patterns and for informing conservation and ecosystem management (Wisz *et al*. 2013). Hutchinson’s niche concept (1957), particularly the distinction between fundamental and realized niches (i.e. the sets of environments where intrinsic population growth rate r_0_ > 0 in the absence and presence of interacting species, respectively), highlights how interspecific interactions reduce population growth in otherwise suitable environments, potentially excluding species from areas where they could persist in the absence of competitors. Differences between fundamental and realized niches quantify the impact of biotic interactions on population dynamics and geographic ranges of species, thereby playing an important role in ecological and biogeographical theory (Louthan *et al*. 2015). However, difficulties in measuring niches, and the fundamental niche in particular, limit quantitative analyses of these concepts - despite their central role in many ecological theories (Laughlin & McGill 2024).

Of particular importance for understanding how realised niches shape geographic distributions and coexistence of species are trade-offs between competitive ability and other life history traits. One long-standing hypothesis is a competition–colonization trade-off that could explains how species can coexist by balancing strong competitive ability with weaker capacity for dispersal and colonization (Tilman 1994, Hastings *et al*. 2025). Additionally, the size of the realized niches of species can be affected by a generalist-specialist trade-off, where generalists can persist across a wide range of environments but get outcompeted by more specialised species under specific conditions (Levins 1968). Generalists and specialists represent two extremes on a spectrum of ecological strategies, each with distinct advantages and trade-offs. Generalists are species that can thrive in a wide range of environmental conditions, whereas specialists are more adapted to specific environmental conditions, often displaying narrow habitat requirements. In fluctuating or unpredictable environments, generalists are more likely to persist because they are not tightly bound to a single resource or habitat. This broader resource use also allows generalists to capitalize on opportunities in diverse conditions, giving them a competitive edge when resources become scarce or unpredictable. As a result, generalists tend to be more resilient to environmental changes, such as shifts in climate or disturbances like droughts or fires. This adaptive flexibility can allow generalists to spread across larger geographical areas, potentially making them more widespread than specialists. However, this ecological flexibility comes with costs, as generalists may not be as efficient at utilizing specific resources as specialists, which can limit their competitive ability in environments within the fundamental niche of specialist species. Specialists can also have a higher population growth rate under benign conditions, further increasing their competitiveness in such environments. The generalist-specialist trade-off can thus promote large-scale species coexistence, where the ability of generalists to use a variety of resources allows them to persist under more conditions, while specialists can thrive in environments where they are highly adapted to specific conditions.

Quantifying fundamental and realized niches at broad scales remains challenging. Species distribution models rarely separate abiotic constraints from biotic interactions (Wisz *et al*. 2013), and geographic patterns alone cannot reliably reveal underlying ecological processes, which may reflect historical legacies and dispersal limits (Dormann *et al*. 2018). Experimental and ecophysiological approaches are also limited in scope and assumptions (Holt 2009; Briscoe *et al*. 2019). In consequence, major gaps remain in our understanding of how biotic interactions shape ecological niches and species’ geographic ranges. In order to close these gaps and, in particular, to understand the relevance of trade-offs between competitive ability and other life-history traits for species ranges, we here quantified fundamental and realized niches from extensive demographic data for 29 closely related plant species. Our study species are shrubs of the Proteaceae family endemic to the Cape Floristic Region, a global biodiversity hot spot (Myers *et al*. 2000). All study species are serotinous: they store their seeds over multiple years in a canopy seedbank until fire triggers seed release, wind-driven seed dispersal, and subsequent establishment of new recruits (Rebelo 2001). Fire is also the predominant cause of mortality of established adults (Bond & Wilgen 1996). This fire-linked life cycle allows the efficient measurement of key demographic rates in single visits to each population (Treuernicht *et al*. 2016) and enabled us to collect data that are informative of variation in population growth rates across each species’ geographic range. In total we combined 5085 population-level measurements of fecundity, recruitment and fire survival across the geographic ranges of the 29 study species (Fig. 1a) with data on the abiotic environmental variation to develop demographic niche models (following Pagel *et al*. 2020, Fig. 1b). We also recorded the population densities of all co-occurring Proteaceae species on all study sites and extended the demographic niche models by effects of heterospecific densities to estimate pairwise competitive effects between our study species.

**Figure 1.**
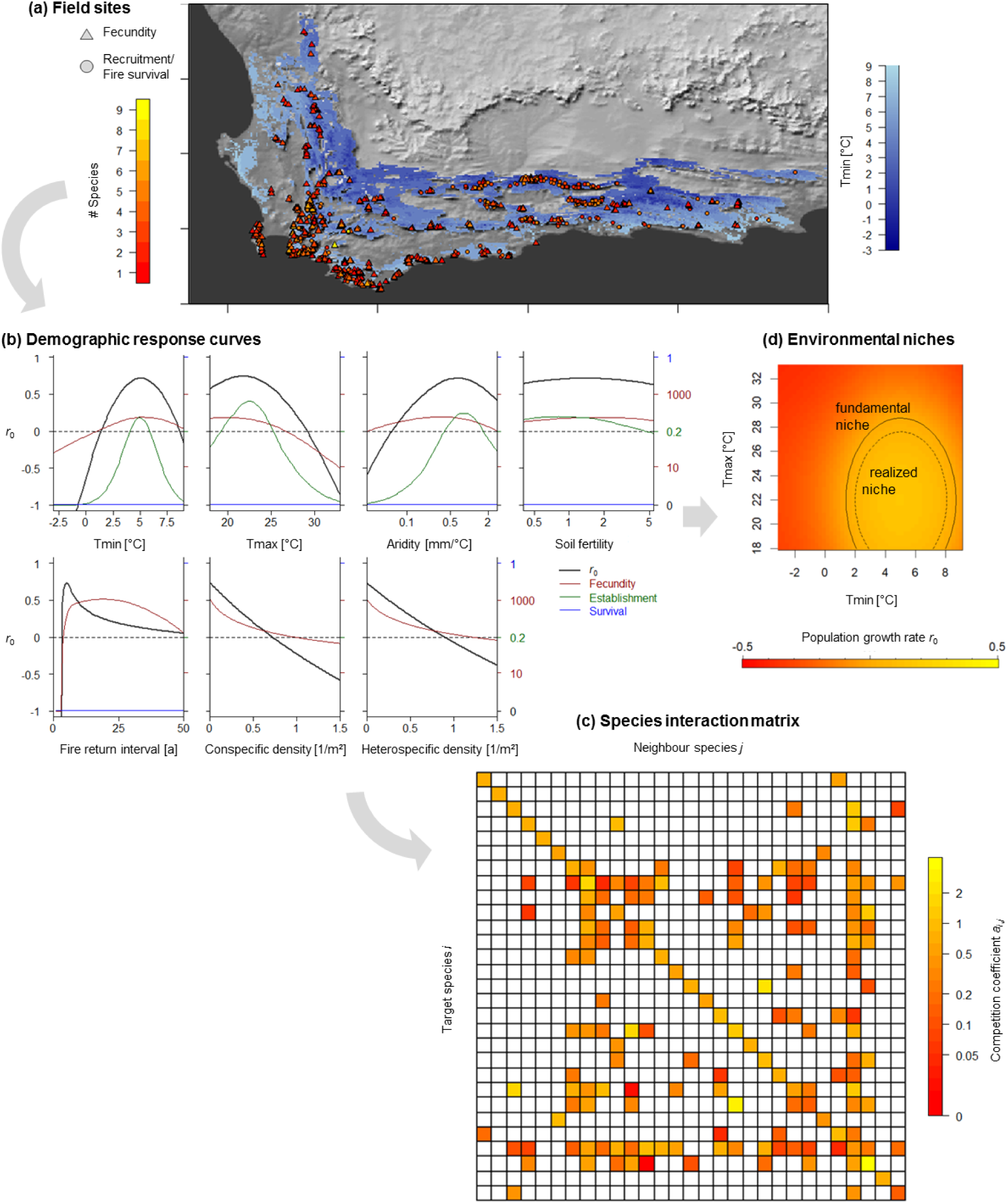
Estimation of demographic response curves, the species interaction matrix and resulting fundamental and realized niches. (a) Geographic distribution of the field sites where key demographic rates were measured across gradients of the abiotic environment and variation in the co-occurrence and abundance of other Proteaceae species. (b) Estimated responses of key demographic rates and the resulting annual intrinsic population growth rate (*r*_0_) of *Leucadendron rubrum* to variation in minimum winter temperature (T_min_), maximum summer temperature (T_max_), indices of summer aridity and soil fertility, fire return interval and the population densities of con- and heterospecific Proteaceae. (c) Matrix of pairwise competition coefficients *a_i,j_*calculated as the effect of the population density of neighbour species *j* on the population growth rate *r*_0_ of target species *i*. (d) 2-d cross-section of the fundamental and realized environmental niche of *Leucadendron rubrum* where *r*_0_ is predicted to be positive either in the absence or presence of interspecific competition.

## Results and Discussion

### Demographic effects of interspecific competition

Including interspecific competition in the demographic models markedly improved explanatory power for both fecundity and recruitment. Specifically, average model fit for fecundity increased from R² = 0.58 to 0.63 (ΔR² = 0.06), while recruitment models showed an even larger gain, with average R² increasing from 0.42 to 0.61 (ΔR² = 0.19). Pairwise effects of heterospecific population densities were estimated individually for each demographic rate and then combined to predict the resulting competitive effect on intrinsic population growth rates (*r*_0_) and to construct a pairwise species interaction matrix of competition coefficients (Fig. 1c).

Species coexistence in local communities is stabilized if the per-capita competitive effect of a species on its own population growth is stronger than the corresponding effects on other species (Chesson 2000). In partial support, we found that for 21 species (73%), intraspecific competitive effects are stronger than the average pairwise effects on other species (mean ratio = 2.58, median = 1.87) (Fig. 1c). For 38 of 67 species pairs (57%), we furthermore predict stable pairwise coexistence, since both species suppress their own population growth more than that of the other species. Moreover, 73% of species showed stronger competitive responses to conspecific neighbours than to heterospecific neighbours also in 73% of cases (mean ratio = 2.61, median = 1.71). Hence, our analysis identifies a predominance of intra-over interspecific competition as one important mechanism stabilizing species coexistence in one of the world’s hottest hotspots of plant diversity (Linder 2003).

### Niche similarity and architecture

From the demographic response curves that describe how each demographic rate varies across environmental gradients (Fig. 1b) we can predict the response of the intrinsic population growth rate (*r*_0_) to key environmental variables (minimum winter temperature, maximum summer temperature, aridity and soil fertility) and thereby quantify species ecological niches as the set of environmental conditions where *r*_0_ is positive in a 4-dimensional hypervolume (Fig. 1d). By setting the population density of other species to zero for these predictions, we specifically quantify the fundamental niche, i.e. the set of environmental conditions where each species is expected to be able to persist in the absence of competitors. Average overall niche similarity between realized niches decrease by 4.4 % (Schöner’s D) compared to fundamental niches. However, the architecture of fundamental niches in relation to the realzed niches (Laughlin & McGill 2024) differed when investigated for different environmnental axes. Only for the minimum winter temperature (T_min_) are the optima of the realized niches more divergent than the optima of the realized niches (by an increase of 6%) and also other niche metrics indicate a centrifugal organization of realized niches along the T_min_-axes, whereas other environmental axis showed characteristics of distinct or shared preferences.

### Generalist-specialist trade-offs

Calculating the sizes of fundamental niches (as the proportion of the environmental space that is covered by the niche) reveals a strong variation across our study species and thus a continuum from environmental specialists to environmental generalists. The average fundamental niches size is 0.381 (median = 0.227) with an standard deviation of 0.352, where pronounced environmental specialists like the dune conebush (*Leucadendron coniferum*) and the rough-leaf conebush (*Leucadendron modestum*) have fundamental niches sizes below 1% (0.00998 and 0.00377, respectively), whereas pronounced generalist like the common sugarbush (*Protea repens*) and the grey-leaf sugarbush (*Protea laurifolia*) include in their fundamental niches the entire range of environmental variation found within the study area.

In order to test the hypothesized generalist specialist trade-off, we regressed the average experienced effects of interspecific competition (relative to the species-specific effect of intraspecific competition) for all study species against their fundamental niche sizes (i.e. the specialist-generalist continuum) and found indeed that interspecific competition acts stronger on generalist species with larger fundamental niches (slope = 0.93, F_1,27_ = 5.534, p = 0.026, adj. R² = 0.139, Fig. 2a). To test the competition–colonization trade-off we regressed the relative strength of interspecific competition against species dispersal ability (Schur *et al*. 2007), but found no significant relationship (slope = 0.529, F_1,27_ = 1.906, p = 0.179, adj. R² = 0.031, Fig. 2c). However, similar to previous analyses of species’ realised niches (Pagel *et al*. 2020) we also found a significant positive effect of fundamental niche size on dispersal ability (slope = 0.486, F_1,27_ = 5.534, p = 0.016, adj. R² = 0.167, Fig. 2b). Such a positive correlation can be expected, if environmental specialists experience selection for lower dispersal distances due to limited availability of suitable habitat (Thompson *et al*. 1999, Pagel *et al*. 2020). Together with the found positive relationship between fundamental niche size and the strength of interspecific competition this could obscure the relationship between dispersal ability and competition strength.

**Figure 2.**
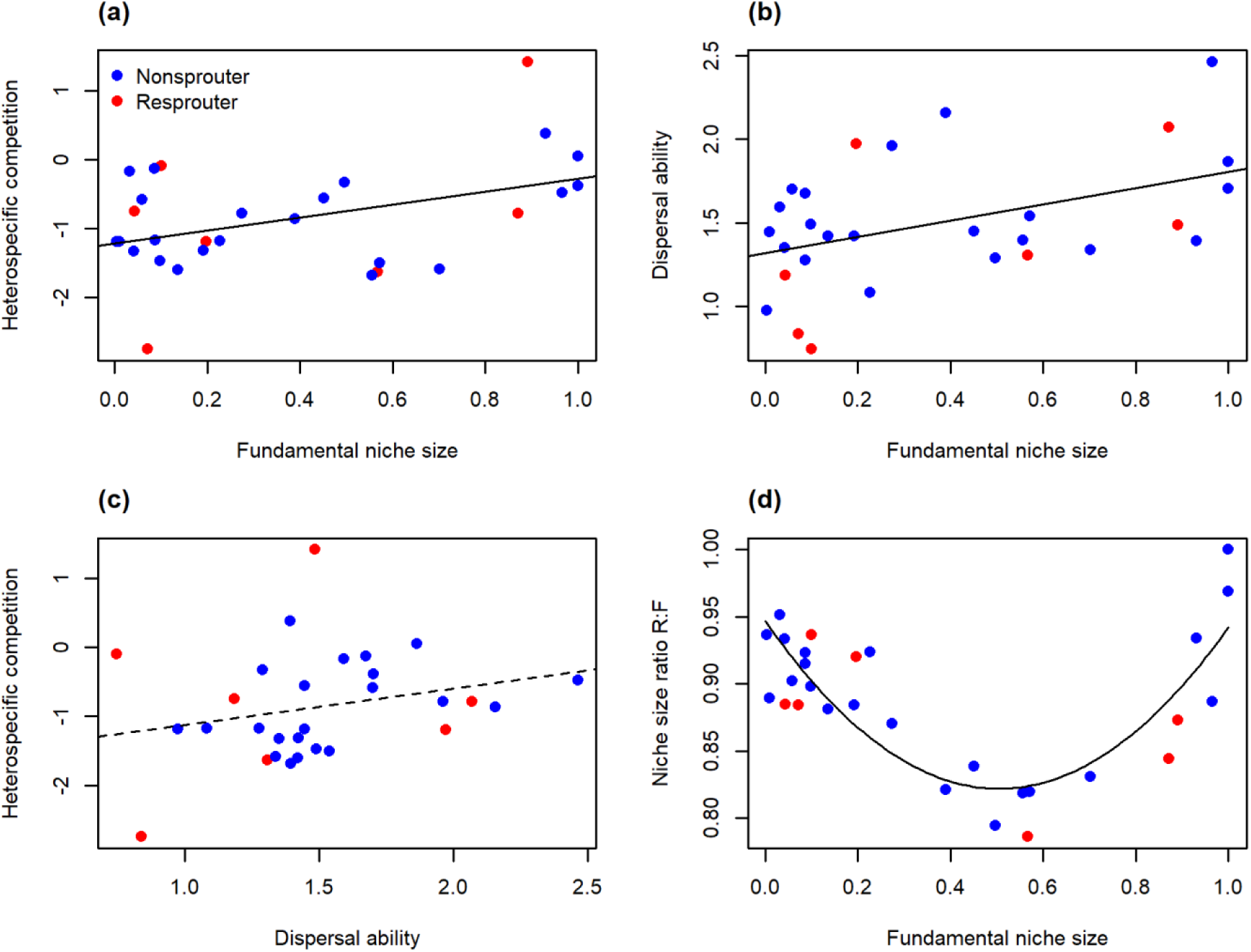
(a) Relationship between species’ normalized fundamental niche size on the relative strength of the effect of heterospecific competition, calculated as *log(average interspecific competition coefficient / intraspecific competition coefficient)* (slope = 0.93, F_1,27_ = 5.534, p = 0.026). (b) Relationship between species’ fundamental niche size and their long-distance dispersal ability (slope = 0.486, F_1,27_ = 5.534, p = 0.016). (c) Relationship between species’ long-distance dispersal ability and the relative strength of the effect of heterospecific competition (n.s.). (d) Relationship between species’ normalized fundamental niche size and the niche size ration *R:F = size of realized niche / size of fundamental niche* (F_2,26_ = 26.425, p < 0.001).

In addition to the estimated strength of interspecific competition, we also analysed the resulting reduction of species’ realized niches relative to their fundamental niches. The sizes of realized niches were estimated from a model that ignored interspecific competition (Pagel *et al*. 2020) and thus predicts where a species is expected to persist under an average degree of competition with other species. As expected, the realized niches are smaller than the respective fundamental niches for all our study species, with an average reduction of the realised niche relative to fundamental niche (1 – R:F, with niche size ration *R:F = size of realized niche / size of fundamental niche*) of 11.2 % (median = 11.3%, min = 0, max = 21.4%). The reduction of the realized niche is small for both extreme specialist and extreme generalist and largest for species with intermediate fundamental niche size (F_2,26_ = 26.425, p < 0.001, adj. R² = 0.645, Fig. 2d). While specialists suffer little from interspecific competition (see above), the small niche differences for extreme generalists emerge, because here both the fundamental and the realized niche span the entire environmental gradient found in our study area. Overall the found differences between fundamental and realized niches are small in comparison to other studies based on occurrence data (Laughlin & McGill 2024). Notably such studies do not represent the full demographic life cycle and ignore mismatches between the realised niche and the actual geographic distribution (Pagel *et al*. 2020, Sandel *et al*. 2025).

### Effects of interspecific competition on potential geographic distributions

Projecting niches into geographic space allows us to also predict potential geographic ranges, and as for the ecological niches we can differentiating a potential fundamental range, where we expect the species to be able to persist in the absence of competition, and a potential realised range, where we expect the species to be able to persist in the presence of competition with other species. Although there are large mismatches between potential and actual ranges of our study species, in particular due to migration limitation (Pagel *et. al.* 2000), studying differences between fundamental a realised potential ranges is important for understanding not only present geographic distributions of species, but also their ability to respond to expected future climate change. Similar to the reduction of species’ realised niches, we find that the potential realised range is reduced compared to the potential fundamental range, although to a slightly smaller degree of 10.6% on average (median = 9.3%, min = 0.7%, max = 21.7%). The proportional reduction of the potential realised range size increases (marginally significant) with potential fundamental range size (slope = 0.0157, F_1,27_ = 3.077, p = 0.091, adj. R² = 0.069), which again highlights that wide-spread generalists tend to be stronger affected by competition.

Potential geographic range can not only be predicted under present conditions, but also for scenarios of future climate change. For a moderate scenario of climate change (ssp370) until the year 2100 we find that both fundamental and realized ranges are predicted to shrink for most our study species (Fig. 3), also there are a few species for which potential geographic ranges could increase. The predicted change of potential realised range sizes under future climate change is negatively related to the current fundamental range size (slope = -0.037, F_1,27_ = 4.571, p = 0.042, adj. R² = 0.113, Fig. 3), whereas such a relationship cannot be found for the change of potential fundamental range sizes (F_1,27_ = 1.831, p = 0.187). Therefore, the loss of future potential range area appears to be aggravated by effects of competition in particular for currently wide-spread generalist species.

**Figure 3.**
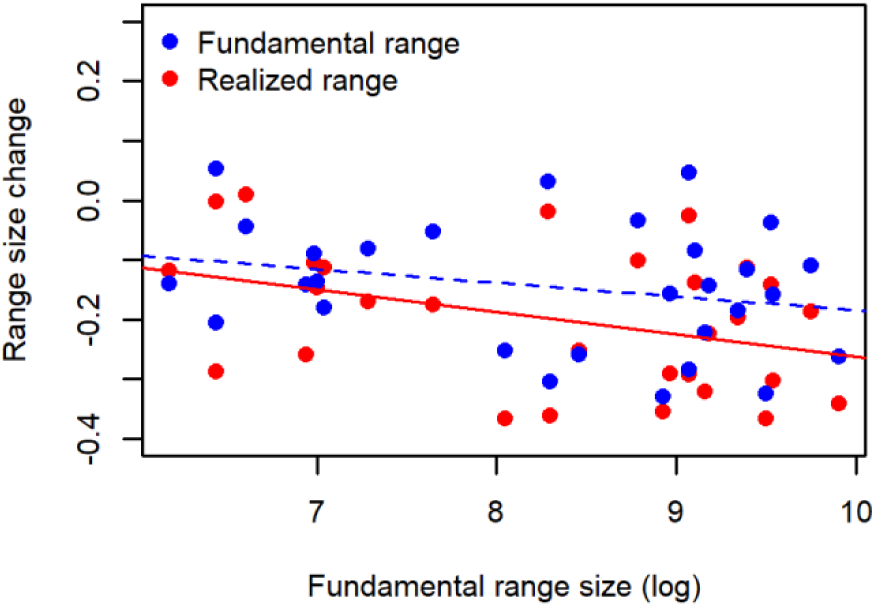
Relationship between the extents of species’ current potential fundamental ranges (area where *r*_0_ > 0 is predicted in the absence of interspecific competition) and the change in either potential fundamental range size or in potential realized range size (area where *r*_0_ > 0 is predicted in the presence of interspecific competition) under a scenario of future climate change until 2100 (fundamental range size (blue line): n.s.; realized range size (red line): slope = -0.037, F_1,27_ = 4.571, p = 0.042).

## Conclusion

In summary, our analyses demonstrate the possibility to quantify both fundamental and realised niches and the basis of observed variation of key demographic rates. The comparison of conspecific and heterospecific density effects on these demographic rates indicates not only, that generally intraspecific competition is stronger than interspecific competition (as one key mechanism for coexistence), but also that generalist species with a wider fundamental niche are stronger affected by interspecific competition than environmental specialists. This generalist-specialist trade-off, which also manifest in larger differences between fundamental and realized niches for generalists, furthermore contributes to the maintenance of Proteaceae biodiversity. However, we did not find evidence for a competition-colonization trade-off, i.e. stronger competitive effects on well dispersed species. This can partly be explained by co-selection of environmental generalism and long-distance-dispersal. But it is also important to consider, that species colonization ability is not only determined by dispersal, but also the capability to establish. It would therefore be important to also study other factor of colonization ability, e.g. seed size and nutrient content (Treuernicht *et al*. 2020). A trait-based understanding also of interspecific interactions would be furthermore relevant for characterizing interaction currencies and to predict biotic interactions in novel communities (Kissling *et al*. 2012).

The found differences between fundamental and realized niches also shape future range dynamics because specialist species, which occupy narrower niches, tend to have smaller geographic ranges and higher extinction risk (see also Treurnicht *et al*. 2021), making them more vulnerable to environmental change. In contrast, generalists often disperse more effectively and more fully occupy their potential ranges (Lester *et al*. 2007, Pagel *et al*. 2020), increasing their capacity to expand under changing conditions. However, actual spread rates depend not only on niche breadth but also on demographic factors such as the intrinsic population growth rate (Svenning *et al*. 2014, Grainger *et al*. 2019) and its reductions due to interspecific competition, so that also generalists might struggle to track shifts of their potential geographic ranges. Our findings thus highlight that the quantification of demographic effects of interspecific interactions on species niches is not only crucial for understanding current biodiversity patterns but also for predicting and mitigating consequences of global change.

## Materials and Methods

### Study Species, Community Composition and Demographic Data

We studied 29 species of the Proteaceae family, specifically of the genera *Protea* (19 species) and *Leucadendron* (10 species). All study species are endemic to the Suth African Fynbos biome (Rebelo 2001). They were chosen to represent variation in geographic ranges as well as variation in functional traits, in particular dispersal and resprouting ability, which have been found to play an important role in shaping species’ niches (Pagel *et al*. 2020). For each species, we obtained data on between-population variation in key demographic rates across the entire life cycle, namely, the total fecundity of adult plants since the last fire (size of individual canopy seed banks), per capita post-fire seedling recruitment (ratio between post-fire recruits and pre-fire adults), and adult fire survival (following the protocols described in Treurnicht et al. 2016). Furthermore, we characterized the community composition on each study site by measuring the population densities of all co-occurring Proteaceae species. For each study species, demographic sampling sites were selected to cover both major environmental gradients across the species’ global geographic distribution and variation in conspecific and heterospecific population densities (Fig. 1a). The final dataset comprised 5,085 population-level records from an average of 183 (median 127) study sites per species.

### Study region and environmental variables

Our study region was defined on a regular grid with a spatial resolution of 1′ × 1′ (c. 1.55 km × 1.85 km) and included all grid cells in which > 5% of the area is covered by Fynbos vegetation (South African National Biodiversity Institute 2012). Climatic and edaphic variables that were previously found to be main determinants of the performance and survival of serotinous Proteaceae (Pagel *et al*. 2020) were extracted from the South African Atlas of Climatology and Agrohydrology (Schulze 2007). We included *January maximum daily temperature* (T_max_), *July minimum daily temperature* (T_min_) and a *January aridity index* (AI) calculated as the ratio between the mean values of precipitation (P) and temperature (T): AI = P/(T + 10°C) (De Martonne 1926). Climatic variables are averages over the years 1950–2000. As an edaphic variable we used a *soil fertility* index that combines soil texture and base status and ranges from 0 to 10 (Schulze 2007). Information on the fire return interval was obtained from both observational records and model predictions. For the demographic sampling sites, information on the fire history (time since the last fire and length of the previous fire interval) was inferred from a combination of measured plant ages, historical records and MODIS satellite observations (see Pagel *et al*. 2020). For predictions of population growth rates across the study region (see below), we used probability distributions of fire return intervals predicted from a climate-driven model of post-fire ecosystem recovery (Wilson et al. 2015).

### Demographic Niche Models

We use three different versions of a demographic response model for the estimation of demographic responses and ecological niches that differ in their representation of interspecific competition. The model #1 does not include any effects of heterospecific population densities on demographic rates (following Pagel et al. 2020). Therefore predictions from this model imply the presence of competing species (specifically the expected biotic environment for any given abiotic environment) and thus predict the realized niches of species. We then extended this model by effects of heterospecific population densities on fecundity and recruitment. In extended model versions, population densities of all co-occurring serotinous overstorey Proteaceae were included as additional predictors of variation in fecundity and recruitment. For model #2, all heterospecific population densities were summed up to estimate a general interspecific density effect. In a final Model #3, we included species-specific effects of the densities of neighbouring species in order to quantify species-specific pairwise competition strength. In the following we first describe the model #1 without effects of heterospecific population densities, as presented in Pagel *et al*. 2020:

#### Demographic response model

We used a hierarchical Bayesian modelling approach for estimating the species-specific responses of key demographic rates (fecundity, per-seed establishment and adult fire survival) to environmental covariates. The model considers effects of climatic and edaphic conditions, variable fire return intervals and intraspecific density dependence at both the adult and the seedling stage. Below, we describe the submodels for variation in each demographic rate.

##### Fecundity

The recorded size of the canopy seed bank (*Seed.count_i,j_*) of plant *j* in population *i* is described by an overdispersed Poisson distribution

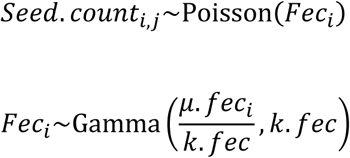

where the expected value of mean fecundity *μ.fec_i_* is determined by limiting effects of post-fire stand age (*Age*), environmental covariates (**X**) and population density(*D*):

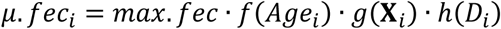

Effects of stand age on fecundity arise from the time of maturation until the first flowering and cone production, increasing accumulation of standing cones on growing plants, cone loss and possibly senescence of aged individuals:

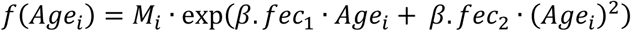

where *M*_i_ is a binary random variable (0, 1) indicating maturity. The probability of population-level maturity is calculated from a Weibull distribution for the age (*t.mat*) of first cone production:

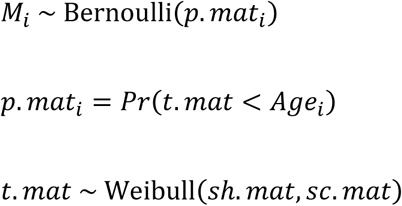

The species-specific time to reproductive maturity (*t.mat*) was constrained to be at least three years for nonsprouters (Rebelo 2001). The effects of the environmental covariates *k* = 1…*K* are described by Gaussian demographic response functions:

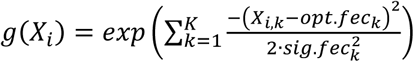

where *opt.fec_k_* denotes the optimal conditions and *sig.fec_k_* measures the width of the response curve. Effects of population density *D_i_* on fecundity are described as (Nottebrock et al. 2017):

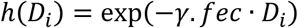

##### Establishment

The establishment of new recruits from seeds is modelled as a binomial process where the number of recruits (#*Recruits_i_*) in population *i* depends on the total number of available seeds (#*Seeds_i_*) in the canopy seed bank at the time of the last fire and the per-seed establishment rate π.*est_i_*:

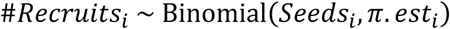

Since #*Seeds_i_*is unknown for recently burned sites where recruitment was recorded, it is modelled as a latent state variable

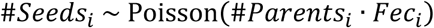

where #*Parents_i_* denotes the number of pre-fire seed sources (only females for dioecious *Leucadendron* species) and *Fec_i_* depends on environmental covariates (**X***_i_*), the post-fire stand age (*Age_i_*) and the adult population density (*D_i_*) at the time of the previous fire as described in the fecundity submodel. Establishment rate π.*est_i_* is affected by environmental covariates (**X***_i_*) and by the densities of seeds *SD_i_* = #*Seeds_i_*/*Area_i_*and fire-surviving adults *AD_i_* = #*Adults_i_*/*Area_i_*.

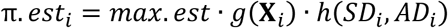

As for fecundity, the effects of different environmental covariates *k* = 1…*K* are described by Gaussian demographic response functions:

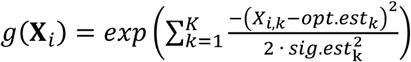

Density effects on establishment result from the density of seeds (*SD_i_*) as well of from the density of fire-surviving adults (*AD_i_*), with different strengths (γ.*est.SD* resp. γ.*est.AD*) for each of these density effects:

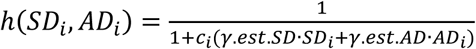

Density-dependent mortality of recruits (self-thinning) is a continuous process, which for Fynbos Proteaceae generally occurs within the first three years after a fire (Manders & Smith 1992). This is described by weighting the density effects with a factor *c*_i_ that depends on the post-fire stand age (*pf.Age_i_*) at the time of sampling:

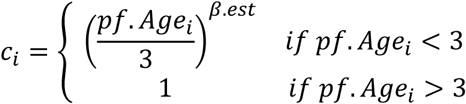

Thereby the model accounts for the fact that more seedlings can be observed if a site is surveyed just shortly after germination (Treuernicht et al. 2016).

##### Survival

Adult fire survival is modelled as a binomial process for the proportion of survivors among all pre-fire adults (#*All.Adults_i_*):

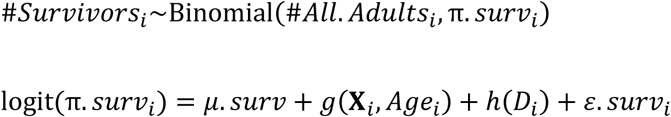

Similar as for fecundity, the effects of different environmental covariates *k* = 1…*K* and of post-fire stand age (*Age_i_*) are described by Gaussian response functions and effects of population density *D_i_* by a negative-exponential function:

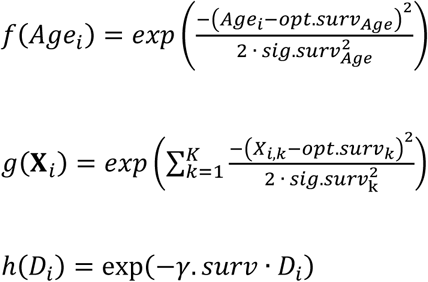

Since adult fire survival rates of nonsprouters are generally low with little intraspecific variation (Treuernicht et al. 2016), we modelled them as species-specific constants and considered effects of covariates only for the survival rates of resprouters.

#### Description of interspecific competition

The previously described model #1 was then extended to include also density effects of all heterospecific Proteaceae on fecundity and recruitment rates. For the fecundity submodel the description of density-dependence was therefore expanded to include effects of not only the local adult density *D*_i,j_ of conspecifics (*j* = *target species*) but also of other species (*j* ≠ *target species*):

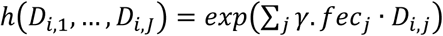

For the recruitment submodel the same expansion was applied both to post-fire adult densities (*AD_i,j_*) and the density of seedlings (*SD_i,j_*). For model #2, population densities were summed over all heterospecifics to estimate general parameters of interspecific competition (γ.*fec_het_*, γ.*est.SD_het_*, γ.*est.AD_het_*). For model #3 species-specific pairwise density effects were estimated when the species co-occurred on a minimum of 10 study sites for each fecundity and recruitment data, and the respective species-specific densities were included. Densities of all other Proteaceae species were summed and combined density effects of all these additional other species were estimated, similar as in model #2. Given the small observed intraspecific variation in fire survival (Pagel *et al*. 2020), no effects of heterospecific population densities on fire survival rates were included in the models.

#### Bayesian parameter estimation and model evaluation

Parameters of the models were estimated independently for each study species. All environmental variables were scaled and centred and the aridity index and soil fertility index were additionally log-transformed before the analyses. The hierarchical model was formulated in a Bayesian framework and samples from the parameter posterior distribution were generated with Markov chain Monte Carlo (MCMC) methods in the software JAGS (Plummer 2003) using three independent MCMC chains with 100,000 iterations after a burn-in period of 500,000 iterations. For each species, we assessed the model fit (using posterior medians) separately for each observed demographic variable (fecundity, recruit:parent ratio, adult fire survival) by calculating Nagelkerke’s general R_N_² (Nagelkerke 1991) relative to null models in which demographic rates (*π.est*, *π.surv*, *μ.fec*) are species-specific constants.

### Quantification of fundamental and realized niches as well as potential ranges

Ecological niches are defined as the set of environmental conditions where the intrinsic population growth rate is positive (*r*_0_ > 0) (Hutchinson 1957). For quantifying species’ realized (post-interactive) niches, *r*_0_ was predicted from model 1 that implicitly includes the effects of other species. For quantifying species’ fundamental (pre-interactive) niches, *r*0 was predicted from the extended model #2 that explicitly estimates the effects of other species and by setting all effects of interspecific competition to zero. Predictions of *r*_0_ were calculated on a 4D grid spanned by the four climatic-edaphic niche axes (_Tmax_, T_min_, AI, and soil fertility). For commensurability of the different niche axes, each environmental variable was scaled by the range of values found across the Fynbos biome, so that each niche axis ranges from zero to one. For *r*_0_ predictions, each scaled niche axis was regularly sampled with a resolution of 0.01 (yielding a hypercube of 10^8^ grid points). Since we focus on species limitation by climatic-edaphic conditions, not disturbance, for each combination of the four climatic-edaphic covariates r_0_ was predicted for the respectively optimal fire return interval. Potential fundamental and realized ranges, respectively, were likewise predicted based on either the model with or without interspecific competition. To geographically project *r*_0_ across the study region, *r*_0_ was not predicted for a single fixed fire return interval but integrated as a weighted geometric mean over the site-specific probability distribution of fire return intervals (Wilson *et al*. 2015, Pagel *et al*. 2020).

### Quantification of pairwise interaction coefficients

Pairwise per-capita density effects *a*_i,j_ of species *j* on species *i* were predicted from the parameters of model #3 (including pairwise density effects on fecundity and recruitment) by a sensitivity analysis of the response of resulting population growth rate *r*_0_ to changes in heterospecific density. For the sensitivity analysis, population growth rates *r*_0_ were predicted at the species-specific environmental optimum while changing the pre-fire population density *D* of either the target species *i* or a single interacting neighbour species *j* from zero to 0.01 m^-1^. Per-capita density effects were then calculated as *a*_i,j_ = Δ*r*_0,i_/ Δ*D_j_*(see Fig. 1c for an overview of estimated pairwise interactions).

### Analyses of relationships between niche characteristics, interspecific competition and dispersal ability

For statistical analyses, the sizes of fundamental and realized niches were calculated as the proportion of the 4D hypercube (see above) spanned by the four standardized climatic-edaphic niche axes (_Tmax_, T_min_, AI, and soil fertility), where population growth rates *r*_0_ are predicted to be positive from the estimated parameters of model #2 and model #1, respectively. Thus, niche sizes are normalized to the ranges of environmental variation found across the Fynbos biome with a maximal possible niche size of one. As a measure of how strongly the realized niche is reduced due to interspecific competition, we the calculated the niche size ration *R:F = size of realized niche / size of fundamental niche* (Laughlin & McGill 2024). Species-specific relative long-distance dispersal ability was derived from a trait-based mechanistic model of primary and secondary wind-dispersal (Schurr et al. 2007) as the number of neighbouring cells on a 1′ × 1′ rectangular grid that can be reached by dispersal from a source cell with a probability of at least 10^-4^. This dispersal measure was then log-transformed and scaled. To analyse species’ responses to climate change, potential fundamental and realized ranges across the Fynbos biome were predicted for a scenario of moderate climate change in 2081-2100 (CMIP6, MIROC6, ssp370, Shiogama *et al*. 2019).

## Data and code availability

Demographic data were partly used under license from CapeNature for the current study, and so are not publicly available. These data are however available from the authors upon reasonable request and with permission of CapeNature. R code and data generated during the analyses is available from the corresponding author upon request.

## Acknowledgements

This work was funded by the German Research Foundation (DFG, grant SCHU 2259/5-2). We are grateful to CapeNature, SANParks, R.M. Cowling and the late W.J. Bond for access to demographic data and to B. Olivier, all field assistants, reserve managers and private landholders who supported our field work. Data were collected under CapeNature permits AAA0028-AAA005-00213, CN35-28-16759 and CN35-28-23767, Eastern Cape Parks permit CRO91/12CR and SANParks permit (Agulhas National Park).

## Author Contributions

J.P. and F.M.S. conceived and designed the study. M.T. collected the data with contributions from F.M.S. J.P. analysed the data and wrote the first draft. All co-authors contributed to the writing of the final manuscript.

